# Evolutionary Diversification of Nitric Oxide Signaling Components Across Metazoa: A Comparative Phylogenomic Analysis

**DOI:** 10.64898/2026.09.15.751659

**Authors:** Ankit Thakur, Mahesh Kulharia

**Affiliations:** Centre for Computational Biology and Bioinformatics, Central University of Himachal Pradesh Dharamshala, District Kangra, Himachal Pradesh - 176215, INDIA

**Keywords:** Nitric Oxide Signaling, Phylogenetics, Gene Duplication, Soluble Guanylate Cyclase, Arginine Metabolism, Metazoan Evolution

## Abstract

Nitric oxide (NO) is an evolutionarily ancient gaseous signaling molecule in animals, yet the evolutionary history of multiple components spanning NO synthesis, substrate regulation, sensing, and signal termination has not been examined in a single integrated phylogenetic framework across Metazoa. Here, we trace the phylogenetic and gene-tree/species-tree histories of ten core NO-pathway components across 89 eukaryotic proteomes spanning Amoebozoa, Excavata, Fungi, Archaeplastida, and Opisthokonta - including Porifera, Placozoa, Cnidaria, Ctenophora, and Bilateria - with a focus on the 69 opisthokont species that anchor the animal comparisons. The results reveal a strikingly modular evolutionary architecture. Nitric oxide synthase (NOS) is broadly conserved across bilaterian lineages, and reconciliation analyses show that NOS diversification was driven predominantly by speciation rather than lineage-specific duplication - supporting the relative conservation of NOS across the sampled bilaterian lineages. By contrast, the arginine-recycling enzymes ASS1 and ASL show ancestral duplications and inferred secondary losses in specific lineages, while arginase isoforms (ARG1/ARG2) and the cGMP-degrading enzyme PDE5A underwent extensive, independent duplications across metazoan groups. GUCY1A1- and GUCY1B1-like sequences were recovered across several metazoan lineages, but the two subunits exhibited partially divergent evolutionary trajectories. Together, these patterns support a model in which a comparatively conserved NO-producing component coexists with more dynamic diversification of associated pathway gene families, providing an evolutionary framework for investigating how NO-cGMP signaling may have been differentially deployed in nervous systems.

## Introduction

Nitric oxide (NO) is a small, membrane-permeant signaling molecule produced in many animal tissues. In the nervous system, NO can act over short distances without vesicular release and is implicated in the regulation of synaptic transmission and plasticity. Evidence for this role is strongest in defined mammalian preparations: in rat hippocampal CA1, pharmacological inhibition of NOS or soluble guanylyl cyclase (sGC) reduces a component of long-term potentiation (LTP), whereas another component is independent of both pathways(Boulton et al., 1995). Studies in rodents have also linked NOS inhibition to altered synaptic plasticity and memory formation(Böhme et al., 1993; Bon and Garthwaite, 2003). These findings support a contribution of NO signaling to selected neural plasticity and behavioral paradigms, rather than a universal requirement for learning or memory.

Mammalian NOS enzymes oxidize L-arginine to generate NO and L-citrulline. In many neurons, activity-dependent Ca2+ entry can activate nNOS (NOS1); eNOS (NOS3) has additionally been detected and functionally implicated in particular hippocampal preparations, but its neuronal expression and contribution should not be generalized across the nervous system(Hopper and Garthwaite, 2006; Kantor et al., 1996). NO activates heme-containing sGC, a heterodimeric receptor that converts GTP to cyclic GMP (cGMP); cGMP then regulates effectors that include cGMP-dependent protein kinases and cyclic-nucleotide-regulated ion channels, while phosphodiesterases hydrolyze cGMP (Boulton et al., 1995)(; Francis et al., 2010). At a subset of CA1 glutamatergic synapses, nNOS localizes postsynaptically and sGCβ presynaptically, consistent with a possible retrograde NO signal; this anatomical arrangement does not establish the same mechanism at all excitatory synapses(Burette et al., 2002)(see Fig. 1).

**Fig. 1.**
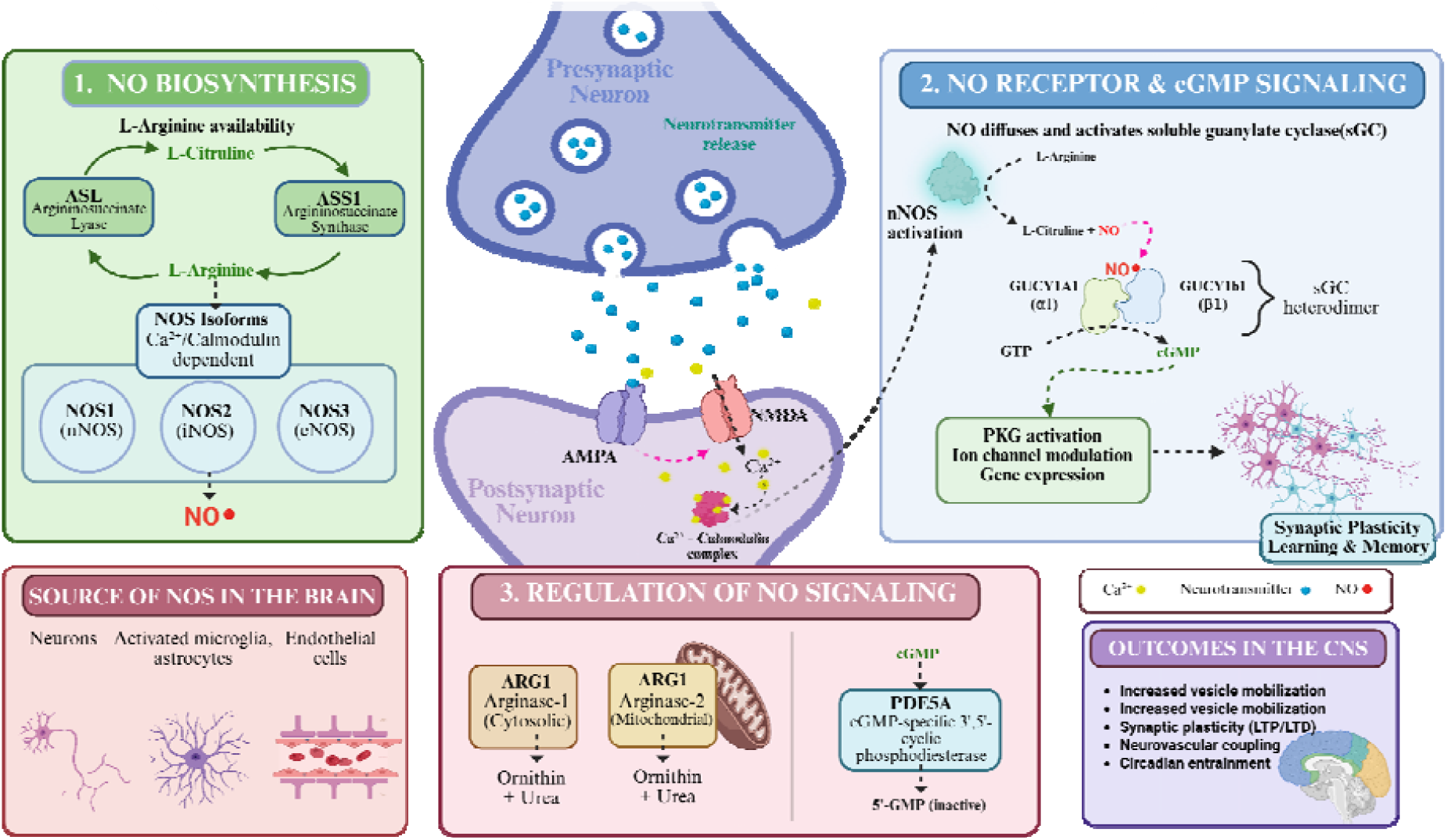
Nitric oxide (NO) signalling in the central nervous system. The schematic illustrates the biosynthesis, signalling cascade, regulation, and physiological roles of nitric oxide (NO) signalling in the central nervous system (CNS). (1) **NO biosynthesis**: Nitric oxide is synthesized from L-arginine by nitric oxide synthase (NOS) isoforms, including neuronal NOS (NOS1/nNOS), inducible NOS (NOS2/iNOS), and endothelial NOS (NOS3/eNOS). Neuronal and endothelial NOS are activated in a calcium/calmodulin-dependent manner. Argininosuccinate synthase (ASS1) and argininosuccinate lyase (ASL) regenerate L-arginine through the citrulline-arginine cycle. (2) **NO-sGC-cGMP signalling**: Following synaptic activation, calcium influx through NMDA receptors activates nNOS in the postsynaptic neuron, leading to NO production. Due to its gaseous and diffusible nature, NO readily diffuses across cellular membranes and activates soluble guanylate cyclase (sGC), composed of GUCY1A1 (α1) and GUCY1B1 (β1) subunits, resulting in cyclic guanosine monophosphate (cGMP) synthesis from GTP. Elevated cGMP activates protein kinase G (PKG), modulating ion channel activity, gene expression, and synaptic plasticity processes associated with learning and memory. (3) **Regulation of NO signalling**: Arginase isoforms ARG1 and ARG2 compete with NOS enzymes for L-arginine metabolism, converting it into ornithine and urea, whereas phosphodiesterase PDE5A terminates signalling by degrading cGMP into inactive 5′-GMP. Major cellular sources of NO in the brain include neurons, activated microglia, endothelial cells, and, under specific conditions, astrocytes. Collectively, NO signalling contributes to neurotransmission, synaptic plasticity, neurovascular coupling, vesicle mobilization, and circadian entrainment.

The availability and compartmentalization of L-arginine provide an additional regulatory layer. Argininosuccinate synthase (ASS1) and argininosuccinate lyase (ASL) can regenerate L-arginine from the NOS coproduct L-citrulline through argininosuccinate. In rat brain, however, ASS and ASL do not uniformly co-localize with one another or with NOS, indicating that local substrate supply can depend on cellular compartmentation and amino-acid transport(Arnt-Ramos et al., 1992; Wiesinger, 2001). Arginase competes with NOS for L-arginine-ARG2 is the mitochondrial isoform described in nervous tissue-and can therefore modulate NO production in a cell- and context-dependent manner(Wiesinger, 2001). Together, enzymes governing arginine availability and catabolism, sGC, cGMP effectors, and phosphodiesterases shape the amplitude, localization, and duration of NO–cGMP signaling.

The evolutionary history of these components remains incompletely resolved. Phylogenetic and syntenic analyses support descent of vertebrate NOS1, NOS2, and NOS3 from a single ancestral invertebrate NOS gene, with recurrent duplication, loss, and domain-architecture changes contributing to present-day NOS diversity(Andreakis et al., 2011a; González-Domenech and Muñoz-Chápuli, 2010a). These studies propose alternative details of duplication timing, so the order of specific events should not be treated as settled. Comparative work has identified NOS homologs in representatives of Porifera, Cnidaria, Placozoa, and Ctenophora, but their retention is uneven: recent ctenophore sampling supports secondary loss in multiple lineages, and sequence detection by itself does not demonstrate catalytic activity or a conserved neural role(Moroz et al., 2020a, 2023). In *Trichoplax*, biochemical and pharmacological evidence supports endogenous NOS activity, whereas studies of ctenophores emphasize that the functional organization of NO signaling remains to be determined(Moroz et al., 2020a, 2023).

Existing evolutionary analyses have concentrated on NOS or on selected NO-receptor components(Andreakis et al., 2011b; Fitzpatrick et al., 2006; González-Domenech and Muñoz-Chápuli, 2010a; Moroz et al., 2023, 2020b). They therefore do not yet resolve the joint history of NOS, sGC subunits, cGMP-metabolizing enzymes, and arginine-supply pathways across a broad sampling of early-branching metazoans. The ctenophore and placozoan comparative studies themselves identify major outstanding functional and comparative questions(Moroz et al., 2020a, 2023). Nor can congruent sequence distribution alone establish that these proteins formed a functionally integrated pathway in ancestral animals. A taxon-rich, component-resolved comparative analysis is needed to distinguish gene presence from duplication and loss, to compare the evolutionary histories of interacting components and identify concordant or divergent patterns, and to identify lineages in which pathway architecture has been retained, modified, or reduced. Here, we investigate the evolutionary distribution and phylogenetic relationships of core NO-signaling and substrate-regulatory components across Metazoa, with emphasis on early-branching animal lineages. This analysis provides a framework for evaluating how the molecular architecture of NO signaling diversified without inferring conserved neural functions from homology alone.

## Methodology

This research investigates the evolution of Nitric Oxide System genes, employing computational methods to analyze proteins and enzymes crucial for Nitric Oxide synthesis, transport, catabolism/regulation, and receptor function across diverse organisms. All proteomic and protein sequence data were obtained from publicly available databases such as NCBI, UniProt-KB, see Supplementary Information A and B). As this study is entirely computational and relies on pre-existing, publicly accessible data, it did not require ethics approval. The methodology followed in this study was taken from *Goulty et al, 2023, Nature communications(Goulty et al., 2023)*, with modifications and updates to suit the objectives of the present study.

### Species Selection

Proteome data for the selected species were primarily collected from publicly available databases such as NCBI and UniProtKB, with additional data sourced from other repositories. To assess the completeness of the proteome data, we utilized the BUSCO tool on 107 proteomes with the eukaryota_odb10 database containing 255 single-copy orthologs. BUSCO estimates the completeness and redundancy of genomic or proteomic data(Seppey et al., 2019). While BUSCO analysis provided valuable insights into the quality of the proteome data, we did not exclude all the species with low BUSCO scores for single copy ortholog, instead we used complete scores and employed a threshold cutoff of ≥ 80. Eventually we ended up with 89 species of proteomes for further study.

### Homology and Orthogroup Identification

We curated a comprehensive set of reference protein sequences; known to be involved in Nitric Oxide synthesis, regulation, and signaling; collectively governing Nitric Oxide homeostasis across peripheral and central nervous tissues, to reconstruct the Nitric Oxide systems across diverse eukaryotic lineages. Based on an extensive literature review and curated biological databases, we included in our study a total of 10 key proteins that represent the core molecular machinery of the Nitric Oxide system. This includes enzymes involved in the synthesis, regulation, and Nitric Oxide dependent signalling receptors.

Annotated protein information was retrieved from the Kyoto Encyclopedia of Genes and Genomes (KEGG)(Kanehisa et al., 2025) database and supplemented with manually curated entries from the literature mining, and UniProtKB/Swiss-Prot database(Pundir et al., 2017; The UniProt Consortium, 2023). These curated reference sequences served as seed queries for homology searches across our dataset of 89 eukaryotic proteomes (69 Opisthokonta species). This approach allowed us to identify potential homologs and reconstruct orthogroups corresponding to Nitric Oxide System-related genes in both well-characterized and poorly annotated species; these served as blast queries sequences for later searches.

We employed PsiBlast(Altschul et al., 1997; McGinnis and Madden, 2004) based searches using the curated query sequences described above. The homologues identified were identified by the PsiBlast (conducted with 3 iterations and an E-value threshold of 1e^−25^) and results retained for downstream analysis (see Supplementary Information C).

The filtered sequences were then subjected BLASTp for functional annotation by querying against the Swiss-Prot database, and the top hit (if sequence coverage >80% and percent identity >40%)(Li et al., 2003; Pearson, 2013; Rost, 1999) from the Swiss-Prot results was used to assign putative functional identity(see Supplementary Information D).

The BLASTP hits across all of the 89 species were sorted by species. The intra-species redundancy was removed using CD-HIT(Huang et al., 2010; Li and Godzik, 2006) with a sequence identity threshold of 100% (-c 1.0) and default parameters (see Supplementary Information E). This step ensured that only unique protein sequences were retained for every species, thereby reducing redundancy in downstream analyses and minimizing redundancy induced computational bias during orthogroup inference.

We employed Broccoli(Derelle et al., 2020), a phylogeny-aware orthology assignment tool that uses a combination of sequence similarity and gene tree-based inference to delineate orthologous groups (OGs) from the filtered protein dataset (see Supplementary Information F). Broccoli was run using default parameters and the maximum likelihood method for gene tree construction, which improves orthology resolution in large, taxonomically diverse datasets.

The resulting OGs of interest i.e., those that potentially represented known components of the Nitric Oxide System system were further analyzed using InterProScan(Jones et al., 2014) (see Supplementary Information G). This combination of automated annotation and expert manual curation ensured accurate functional classification of Nitric Oxide System-related protein families across a wide phylogenetic range.

We grouped all the identified sequences into three main datasets: one for receptors, one for Nitric Oxide synthesis, and one for its regulation. For each of these three datasets, we used the orthogroups identified by Broccoli. Specifically, we selected orthogroups based on their InterProScan domain annotations: those annotated as receptor-like proteins were used for the receptor dataset, proteins “likely to be involved” in Nitric Oxide synthesis for the synthesis dataset, and regulation-like proteins for the regulation enzyme dataset.

To explore sequence space relationships across these three functional groups, we employed CLANS2 version:2.2.2 (CLuster ANalysis of Sequences)(Frickey and Lupas, 2004), a tool that visualizes pairwise sequence similarity in 2D/3D using an all-vs-all BLAST-based clustering approach. Each group of receptors, synthetic enzymes, and regulating enzymes, was analyzed independently. CLANS2 was run with pairwise similarity thresholds (P-values) ranging from 1e^−15^ to 1e^−100^. At more permissive thresholds (e.g., 1e^−20^), clusters appeared larger and more interconnected, capturing distant homologies. As the stringency was increased (e.g., 1e^−100^), clusters resolved into more distinct groups, allowing finer discrimination of protein subfamilies and evolutionary lineages (see Supplementary Information H). Final values of thresholds were empirically determined based on cluster stability across stringency gradients.

Sequences that formed well-defined clusters in CLANS at specific stringency levels were retained for downstream functional and phylogenetic analysis. For receptors, sequences that formed stable and coherent clusters at a P-value threshold of 1e^−37^ were selected, while loosely connected or isolated sequences were omitted.

Similarly, for biosynthetic enzymes, a P-value of 1e^−37^ was found to be optimal for separating functionally coherent clusters. For Regulating enzymes, a threshold of 1e^−49^ was required. To assess the robustness of sequence exclusion decisions we performed repeated clustering by varying threshold values and examined sequences that appeared or disappeared across threshold levels for taxonomic identities.

### Phylogenetic Tree Construction and Analysis

Based on the CLANS clustering results, only sequences that formed coherent and well-supported clusters were retained from each orthogroup (OG) for phylogenetic investigation. These OGs, corresponding to distinct Nitric Oxide System gene families (e.g., biosynthetic enzymes, regulating enzymes, and receptors), were aligned independently using MAFFT(Katoh et al., 2019) (see Supplementary Information I).

Post-alignment, positions with more than 70% gaps were removed using trimAl(Capella-Gutiérrez et al., 2009) to eliminate poorly aligned and phylogenetically uninformative regions. This step was crucial for reducing noise and improving the accuracy of tree reconstruction.

Phylogenetic trees were then reconstructed using IQ-TREE2(Minh et al., 2020), employing ModelFinder to automatically select the best-fit substitution model based on the Bayesian Information Criterion (BIC) and the resulting tree topologies were visualized, colored, and annotated using FigTree v1.4.5 (https://tree.bio.ed.ac.uk/software/figtree/). To assess nodal support, 1,000 ultrafast bootstrap (UFB)(Hoang et al., 2018) replicates were calculated. Additionally, Transfer Bootstrap Expectation (TBE)(Zaharias et al., 2023) values were computed using 100 non-parametric bootstrap replicates to provide more robust support for deep or weakly supported nodes (see Supplementary Information J for iqtree2 computation and Supplementary Information Na/b for coloured tree pdf files).

### Rogue taxa analysis

We employed two complementary methods: the t-index and the Leaf Stability Index (LSI)(Wilkinson, 2006), to identify and eliminate unstable or rogue sequences that could distort phylogenetic inference.

The t-index, implemented in IQ-TREE2, evaluates how frequently a taxon changes its phylogenetic position across bootstrap replicates. High t-index values indicate topological instability, suggesting that the taxon may be a rogue element affecting overall tree resolution. We primarily relied on TBE-based t-index scores, and sequences with a t-index > 2 were classified as unstable.

The Leaf Stability Index (LSI) provides a complementary measure by estimating the frequency with which a taxon maintains a consistent phylogenetic placement across bootstrap trees. It is calculated using quartet frequencies, making it less sensitive to small topological rearrangements. LSI was computed using RogueNaRok(Aberer et al., 2013), with TBE-based trees. In this metric, lower LSI scores denote higher instability.

Taxa flagged as unstable by either method t-index > 2 or low LSI scores were considered problematic. These rogue sequences were excluded from the final phylogenetic trees to prevent topological artifacts that could compromise downstream reconciliation analyses. Pruning was carried out using the rnr-prune function of RogueNaRok (see Supplementary Information K). This combined approach allowed us to refine our gene trees by minimizing the impact of highly unstable taxa, thereby enhancing both tree robustness and biological interpretability.

### Reconciliation analysis

To investigate the evolutionary history of Nitric Oxide System gene families in the context of species evolution, we performed gene-tree::species-tree reconciliation using GeneRax(Morel et al., 2020) (--si-strategy HYBRID -r UndatedDL), an advanced tool designed for inferring duplication, transfer, and loss (DTL) events.

We used the undated duplication-loss (UndatedDL) model (-r UndatedDL), which allows reconciliation of gene trees with a species tree without requiring branch length information in absolute time units. This approach is particularly useful for diverse datasets where molecular dating is either unavailable or unreliable. Maximum likelihood gene trees were generated by IQ-TREE2 for the unrooted gene tree input. Subsequently, rather than inferring a species tree *de novo*, we utilized the Open Tree of Life (OToL)(Hinchliff et al., 2015) to provide a robust phylogenetic framework for our analysis. The comprehensive species tree was pruned to include only the 89 taxa corresponding to the proteomes used in this study. By adopting this externally curated topology, we ensured that the species (see Supplementary Information L) tree was independent of the gene trees generated during our orthology search, thereby eliminating potential circularity in the downstream reconciliation process.

The substitution model for each orthogroup (OG) was specified based on the best-fit model determined during gene tree construction in IQ-TREE2 using the Bayesian Information Criterion (BIC)(Zhao et al., 2008).

The reconciliation output was visualized using ThirdKind(Penel et al., 2022) (see Supplementary Information Ma/b), a tool designed to interpret and graphically represent gene duplication, loss, and transfer events in the context of species evolution. This facilitated clear identification of evolutionary dynamics underlying the distribution and diversification of Nitric Oxide System-related gene families.

### Data Visualization

Schematic illustrations and data visualizations were prepared using standard graphic design tools. Figures 1-4 were created using the vector graphics editor Inkscape (https://inkscape.org/).

## RESULTS

### Synthesis

The nitric oxide (NO)–synthesizing enzymes nitric oxide synthase (NOS), argininosuccinate synthase 1 (ASS1), and argininosuccinate lyase (ASL) were analyzed as key components that increased NO availability. NOS directly catalyzed NO production, whereas ASS1 and ASL supported arginine biosynthesis and thereby supplied the substrate required for NO synthesis(Erez et al., 2011; Marletta et al., 1998; Qualls et al., 2012). We examined these enzymes separately to compare their evolutionary trajectories and determine whether their patterns of diversification were broadly concordant or divergent within the NO-producing pathway.

### Nitric Oxide Synthase (NOS) Evolution

Our analysis identified homologs of NOS across all examined bilaterian lineages, including Acoela, Spiralia, Ecdysozoa, Ambulacraria, and Chordata. Phylogenetic reconstruction showed strong topological support (TBE = 0.99; UFB = 98; see Fig. 1A), indicating high confidence in node stability. Gene tree::Species tree reconciliation further showed that speciation events were the main evolutionary force shaping NOS diversification within this clade, with little evidence for lineage-specific gene duplication or loss.

### The L-Arginine Recycling Pathway: ASS1 and ASL

The distribution of Argininosuccinate Synthase (ASS) spans nearly all Metazoa, including Porifera, Placozoa, and Cnidaria (Anthozoa and Medusozoa), as well as Bilateria. Notably, ASS1 was absent in Ctenophora. While ASS1 candidates were detected in Acoela (*Symsagittifera roscoffensis*) and the outgroup Choanoflagellata (*Salpingoeca rosetta*), these sequences exhibited phylogenetic instability (long-branch attraction or compositional bias) and were excluded from the final reconciliation to maintain tree topology integrity. The ASS1 clade was strongly supported (TBE = 0.99, UFB = 100), and Gene tree::Species tree reconciliation analysis revealed ancestral duplication events, suggesting a more complex paralogous history than that of NOS(see Fig. 2B).

**Fig. 2.**
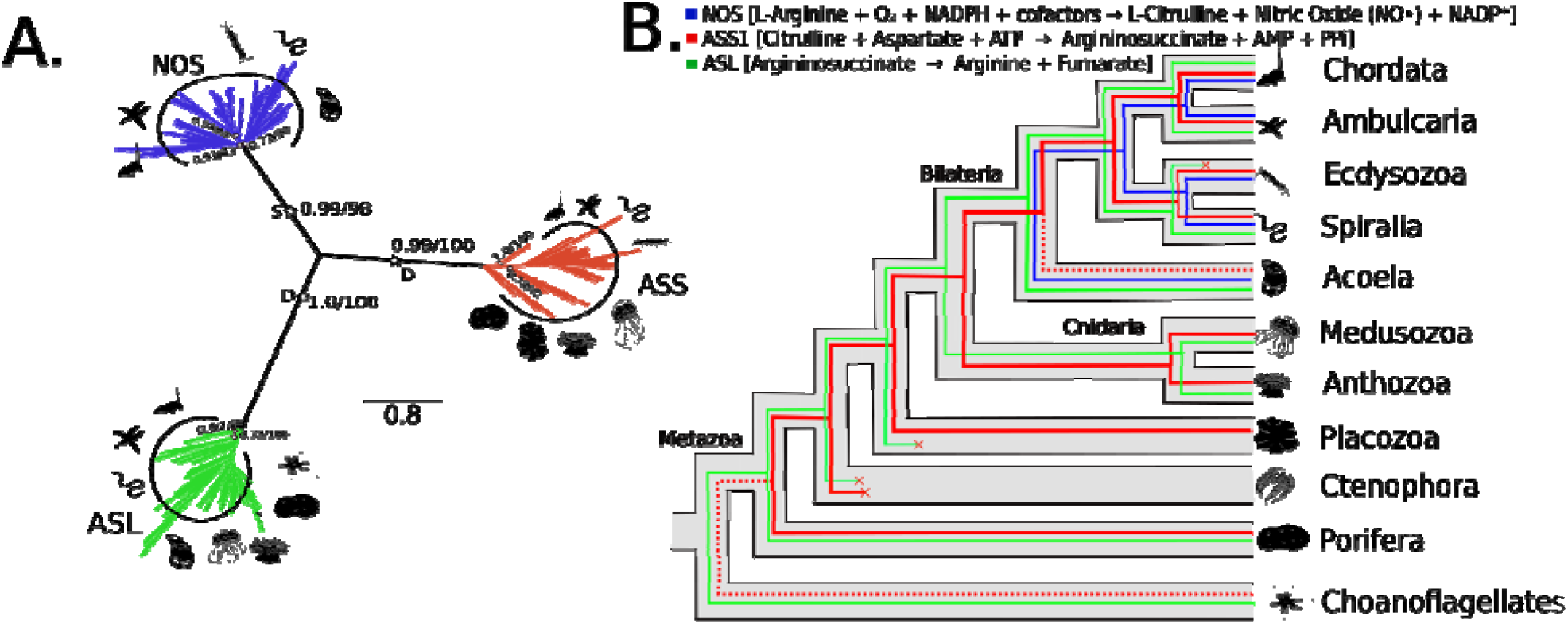
Phylogeny and reconciliation for Nitric Oxide Synthase (NOS), Argininosuccinate Synthase (ASS1) and Argininosuccinate Lyase (ASL). (A) Transfer bootstrap expectation tree and (B) simplified illustration of reconciliation calculated using GeneRax for NOS, ASS1 and ASL sequences. The nodal supports shown are transfer bootstrap expectation (TBE) scores (in decimal values), and ultrafast bootstrap proportion supports (in whole numbers) for key nodes. Star mark(☆, if present) represents the Speciation (S) or Duplication (D) event with TBE and UFB support values. The rectangular box(▄, if present) represents the S or D event of a given branch. The circular shape (○, if present) on nodes represents the TBE/UFB events for particular node. Silhouettes were obtained from Phylopic.org.

Similarly, Argininosuccinate Lyase (ASL) orthologs were recovered across Porifera, Anthozoa, Medusozoa, and several bilaterian lineages (Acoela, Spiralia, Ambulacraria, and Chordata). Conversely, our analysis indicates inferred secondary losses of ASL in Ctenophora, Placozoa, and Ecdysozoa. The resulting ASL clade was recovered with near-unanimous statistical support (TBE = 1.0, UFB = 100). Gene tree::Species tree reconciliation inferred that the ASL topology was significantly shaped by gene duplication events. These duplications indicate increased evolutionary complexity within the ASL-associated gene family and raise the possibility that lineage-specific changes in arginine metabolism contributed to differences in NO-related metabolic capacity.

### Phylogenetic diversification of NO-regulatory components

The nitric oxide (NO) - regulating enzymes phosphodiesterase 5A (PDE5A), arginase-1 (ARG1), and arginase-2 (ARG2) were analyzed as key modulators of NO signaling. PDE5A was responsible for cGMP hydrolysis, thereby controlling the downstream output of the NO-sGC-cGMP pathway, whereas ARG1 and ARG2 competed with nitric oxide synthases for arginine and thereby influenced NO availability(Kass et al., 2007; Que et al., 2002). We examined these enzymes separately to compare their evolutionary trajectories and determine whether their patterns of diversification were broadly concordant or divergent within the NO regulatory network.

### Evolutionary Dynamics of Arginase Isoforms

Phylogenetic analysis of the Arginase family revealed that **Arginase-1 (ARG1)** is widely distributed across Placozoa, Cnidaria (Anthozoa and Medusozoa), and Bilateria but absent in Acoela. This clade demonstrated maximal statistical support (TBE=1.0, UFB=100; see Fig. 3A). Similarly, **Arginase-2 (ARG2)** was identified in Anthozoa and throughout various Bilaterian lineages but absent in Porifera, Ctenophore, Placozoa, Medusozoa and Acoela. Notably, ARG1 and ARG2 sequences did not form reciprocally monophyletic clades; instead, terminal nodes (leaves) were interspersed, suggesting high sequence homology or a series of complex, lineage-specific duplication events rather than a single ancient divergence. Gene tree::Species tree reconciliation confirmed extensive gene duplications within both Arginase lineages (see Fig. 3B).

**Fig. 3.**
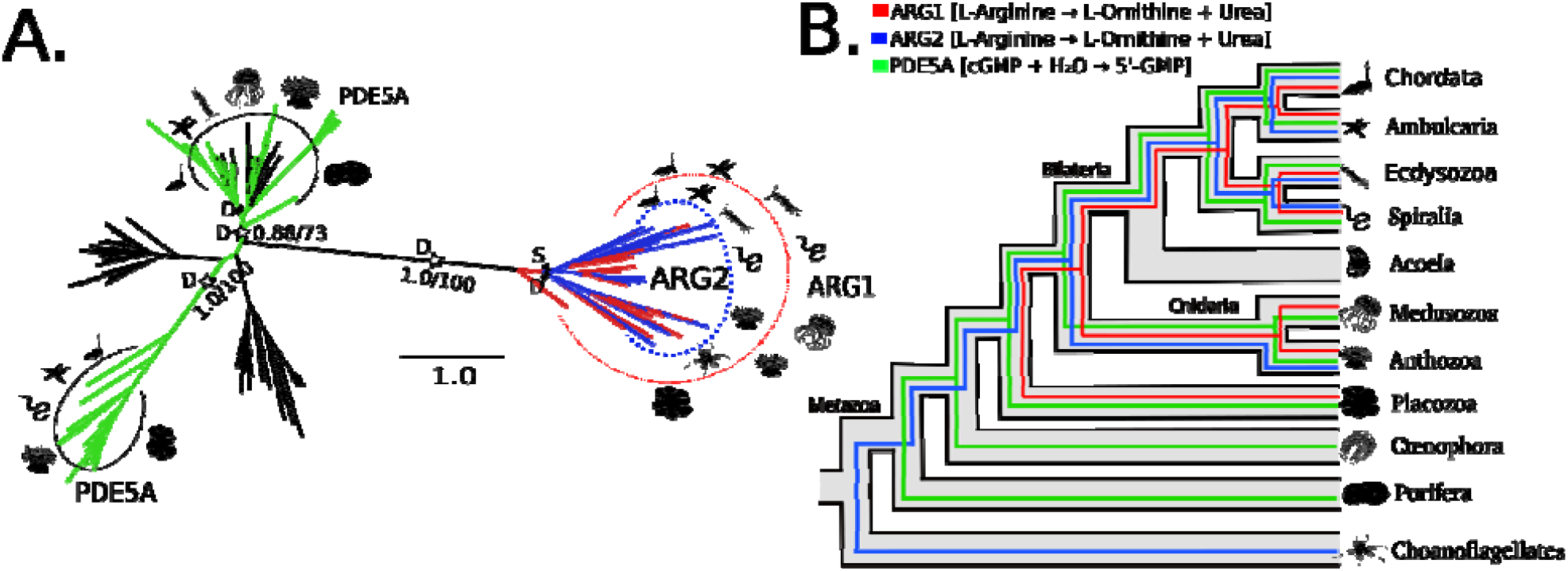
Phylogeny and reconciliation for Arginase-1 (ARG1), Arginase-2 (ARG2) and 3’,5’-cyclic phosphodiesterase (PDE5A). (A) Transfer bootstrap expectation tree and (B) simplified illustration of reconciliation calculated using GeneRax for ARG1, ARG2 and PDE5A sequences. The nodal supports shown are transfer bootstrap expectation (TBE) scores (in decimal values), and ultrafast bootstrap proportion supports (in whole numbers) for key nodes. Star mark(☆, if present) represents the Speciation (S) or Duplication (D) event with TBE and UFB support values. The rectangular box(▄, if present) represents the S or D event of a given branch. The circular shape (○, if present) on nodes represents the TBE/UFB events for particular node. Silhouettes were obtained from Phylopic.org.

### Diversification of Phosphodiesterase 5A (PDE5A)

The cGMP-specific 3’,5’-cyclic phosphodiesterase (**PDE5A**) was recovered across a broad taxonomic range, including Porifera, Ctenophora, Placozoa, Cnidaria (anthozoa and medusozoa), Spiralia, Ecdysozoa, Ambulacraria, and Chordata but absent in Acoela. Our phylogenetic reconstruction resolved two distinct PDE5A clades. The first clade comprised sequences from Placozoa, Cnidaria, and Bilateria, exhibiting robust support (TBE=1.0, UFB=100). The second clade, which included Porifera, Cnidaria, and various Bilaterian representatives, showed moderate support (TBE=0.88, UFB=73). Gene tree::Species tree reconciliation analysis indicates that these two clades likely arose from an ancestral duplication event, with subsequent independent duplications occurring within each lineage.

### Evolution of the Soluble Guanylate Cyclase (sGC) Heterodimer

In established vertebrate systems, soluble guanylate cyclase (sGC) α1 and β1 subunits form a functional heterodimer that senses NO and catalyzes the conversion of GTP to cGMP(Shiga and Suzuki, 2005; Zhou et al., 2004). In this study, GUCY1A1 and GUCY1B1 homologs were analyzed separately to compare their phylogenetic and gene-tree/species-tree reconciliation patterns.

### The Alpha-1 Subunit (GUCY1A1)

The alpha-1 subunit (**GUCY1A1**) exhibited a similar taxonomic distribution, with sequences recovered from Placozoa, Anthozoa, and Bilateria (Spiralia, Ecdysozoa, Ambulacraria, and Chordata) but absent in Medusozoa and Acoela. This clade was recovered with maximal statistical support (TBE = 1.0, UFB = 100; see Fig. 4A). In contrast to the speciation-driven divergence noted in some beta-subunit lineages, Gene tree::Species tree reconciliation of the GUCY1A1 clade primarily indicated a history of gene duplication(see Fig. 4B). These results highlight complex and partially divergent evolutionary trajectories for the GUCY1A1- and GUCY1B1-associated gene families, characterized by inferred speciation and lineage-specific duplication events.

**Fig. 4.**
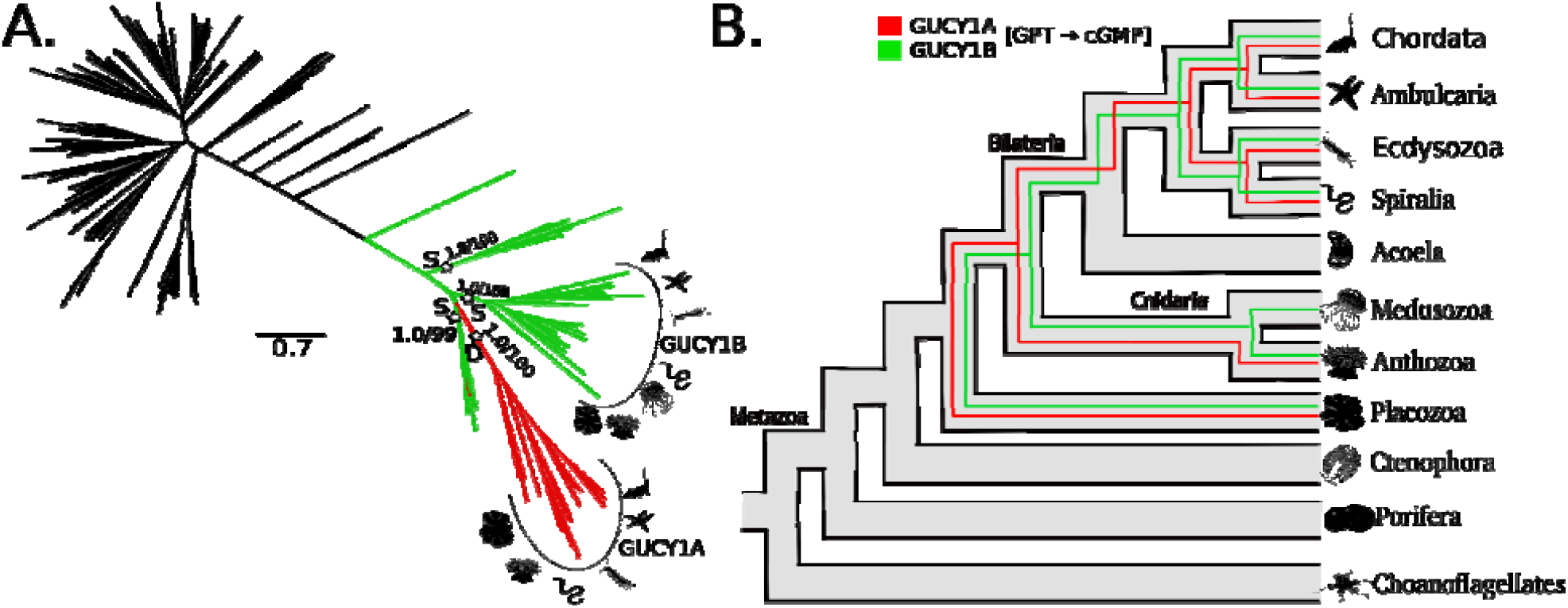
Phylogeny and reconciliation for Alpha-1 Subunit (GUCY1A1) and Beta-1 Subunit (GUCY1B1). (A) Transfer bootstrap expectation tree and (B) simplified illustration of reconciliation calculated using GeneRax for GUCY1A1 and GUCY1B1 sequences. The nodal supports shown are transfer bootstrap expectation (TBE) scores (in decimal values), an ultrafast bootstrap proportion supports (in whole numbers) for key nodes. Star mark(☆, if present) represents the Speciation (S) or Duplication (D) event with TBE and UFB support values. The rectangular box(▄, if present) represents the S or D event of a given branch. The circular shape (○, if present) on nodes represents the TBE/UFB events for particular node. Silhouettes were obtained from Phylopic.org.

### The Beta-1 Subunit (GUCY1B1)

The soluble guanylate cyclase beta-1 subunit (**GUCY1B1**) was identified across a diverse range of Metazoan lineages, including Placozoa, Cnidaria (Anthozoa and Medusozoa), and various Bilaterian clades (Spiralia, Ecdysozoa, Ambulacraria, and Chordata) but absent in Acoela. Phylogenetic analysis recovered a highly supported monophyletic assemblage (TBE = 1.0, UFB = 100; see Fig. 4A). Interestingly, the GUCY1B1 topology revealed a core “main” clade and several divergent lineages (distal leaves) that appear phylogenetically distinct. Gene tree::Specie tree reconciliation indicated ancestral speciation events(see Fig. 4B). This suggests an early diversification of the beta subunit prior to the radiation of major metazoan groups.

## Discussion

Our comparative analysis of ten candidate components of NO signaling across the sampled eukaryotic proteomes recovered a heterogeneous evolutionary pattern rather than a single shared history. The data directly show taxon-specific recovery of putative homologs and gene trees with family-specific reconciliation histories: NOS was broadly recovered among the sampled bilaterians, whereas ASS1/ASL, arginase-family homologs, PDE5A-like homologs, and the putative sGC subunit families showed different inferred patterns of duplication, loss, and lineage distribution. These results are conditional on the sampled proteomes, sequence annotation, orthogroup assignment, gene-tree reconstruction, and reconciliation model. They therefore support inferences about the history of the recovered gene families, but do not by themselves demonstrate that all recovered homologs retain the same enzymatic activity, expression pattern, pathway membership, or physiological role.

For NOS, our analysis recovered homologs in all examined bilaterian lineages and the reconciliation of the retained NOS clade inferred predominantly speciation events, with relatively little lineage-specific duplication or loss in this dataset. This is consistent with predominantly vertical inheritance of the sampled bilaterian NOS homologs. It should not be interpreted as evidence that NOS evolution was uniformly static across Metazoa or that an identical NO-producing function was maintained in every lineage. Previous comparative work found that animal NOS proteins retain broadly conserved domain and gene architectures, while also documenting recurrent duplications and changes in protein architecture in different lineages(Andreakis et al., 2011a) (Andreakis et al., 2011). Likewise, the three mammalian NOS isoforms are inferred to derive from a single ancestral invertebrate NOS gene, whereas labels such as “neuronal” and “inducible” should not be applied to invertebrate NOS homologs solely on phylogenetic grounds(González-Domenech and Muñoz-Chápuli, 2010b).

ASS1 and ASL showed a more discontinuous pattern in the present analysis. ASS1 homologs were recovered across many sampled metazoans but not from the sampled ctenophore proteomes, and the retained ASS1 clade showed inferred ancestral duplications. ASL homologs were recovered from poriferan, cnidarian, and several bilaterian samples, while reconciliation inferred lineage-specific losses and duplications. These findings identify candidate changes in the evolutionary histories of ASS1- and ASL-like genes; they do not establish that a complete citrulline–arginine cycle was present, absent, or rate-limiting for NO production in the corresponding taxa. In characterized mammalian and vascular-cell contexts, ASS and ASL regenerate arginine from citrulline produced during NOS activity and can provide substrate for high-output NO synthesis(Mori and Gotoh, 2000; Xie et al., 2000). Whether the duplicated or retained homologs identified here alter arginine flux, NOS substrate availability, or NO output in non-model metazoans remains an experimental question.

The regulatory families also displayed distinct gene-family histories. In the sampled proteomes, arginase-family homologs assigned as ARG1 or ARG2 showed lineage-specific distributions and extensive inferred duplications; the non-reciprocal monophyly reported for these groups means that isoform assignments and one-to-one functional equivalence should be made cautiously outside well-characterized taxa. PDE5A-like homologs formed two clades, for which the reconciliation inferred an ancestral duplication followed by additional lineage-specific duplications. In mammalian cells, arginase can lower intracellular arginine and thereby modulate

NO production, and PDE5 hydrolyzes cGMP to shape cGMP signaling(Mori and Gotoh, 2000). These established biochemical roles provide a rationale for examining these families alongside NOS and sGC, but the present phylogenetic analysis does not show that duplication changed NO availability, cGMP turnover, or signaling dynamics in any lineage. Direct measurements of substrate specificity, enzyme kinetics, localization, expression, and NO/cGMP dynamics will be required to test those possibilities.

The putative sGC alpha- and beta-subunit families likewise had different inferred histories. GUCY1A1-like sequences were recovered from several sampled placozoan, anthozoan, and bilaterian taxa and were associated primarily with inferred duplications, whereas GUCY1B1-like sequences were recovered from a broader set of sampled cnidarian and bilaterian taxa and showed inferred ancestral speciation events. In mammals, soluble guanylate cyclase is an NO-responsive alpha1beta1 heterodimer: NO binding to the beta1 heme-containing sensor domain triggers conformational changes that increase cyclase activity and cGMP production(Kang et al., 2019). The co-occurrence of alpha- and beta-like homologs in some sampled lineages is consistent with retention of sGC-related genes, but does not demonstrate heterodimerization, NO responsiveness, functional dependence, or co-evolution in those organisms. Establishing co-evolution would require explicit correlated-evolution analyses and, critically, interaction and functional data from representative species.

The phylogenetic results are biologically relevant to neural signaling because experimental work in vertebrate systems has implicated NO/cGMP signaling in particular forms of synaptic plasticity. Reviews and pharmacological experiments support roles for NO in hippocampal LTP, cerebellar LTD, and selected learning and memory paradigms; however, these effects differ across preparations and behavioral tasks(Böhme et al., 1993; Haley and Schuman, 1994). NO is also a short-lived diffusible messenger, a property compatible with local, activity-dependent signaling in characterized nervous systems(Haley and Schuman, 1994). Our analyses neither measured neuronal expression nor tested synaptic, electrophysiological, behavioral, or cognitive phenotypes. Accordingly, they do not show that the observed gene-family histories produced changes in plasticity, learning, memory, neural complexity, or intelligence-related traits.

Several earlier studies provide useful context but do not remove these limits. Andreakis et al. (2011) and González-Domenech and Muñoz-Chápuli (2010) addressed NOS evolution across metazoans and vertebrates, respectively, while Moroz and colleagues documented three distinct NOS genes in placozoans and combined expression and inhibitor-sensitive biochemical measurements to demonstrate functional NOS activity in that lineage(Andreakis et al., 2011a; González-Domenech and Muñoz-Chápuli, 2010b; Moroz et al., 2020a). The latter example illustrates why phylogenetic recovery and functional assignment should remain separate: functional activity was supported there by biochemical perturbation and metabolite measurements, not by gene presence alone. In contrast, the present study broadens the comparison by considering NOS-associated arginine-metabolism genes, putative cGMP-regulatory genes, and putative sGC subunits within a shared taxonomic framework.

Taken together, the results support the hypothesis that homolog families associated with NO synthesis, arginine metabolism, NO sensing, and cGMP turnover did not all follow the same evolutionary trajectory in the sampled taxa. The combination of broad NOS recovery with more variable reconciliation patterns in several other families provides a basis for targeted functional comparisons among lineages. It may be useful to test whether particular duplication or loss events are associated with changes in expression, catalytic properties, protein interactions, or cellular NO/cGMP responses. Such associations, if demonstrated in future work, could clarify how different components of NO-related signaling acquired lineage-specific roles. At present, the results should not be taken to imply that gene duplication enhanced signaling, that gene loss represented adaptation, or that molecular diversification caused increases in neural or cognitive complexity.

Several limitations require emphasis. The analysis is based on available proteomes and on homology, domain annotation, phylogenetic reconstruction, and gene-tree/species-tree reconciliation; apparent absences may therefore reflect incompleteness, annotation error, divergent sequences below detection thresholds, or the filtering steps used, rather than true gene loss. Reconciliation provides model-based inferences of duplication, loss, and speciation conditional on the inferred gene and species trees; it cannot independently establish the biochemical identity or physiological role of a sequence. In particular, the absence of direct assays prevents assignment of NOS activity, arginine recycling, arginase competition, PDE5A-like cGMP hydrolysis, or sGC heterodimerization to the recovered homologs. Developmental and tissue-resolved expression studies, biochemical characterization, interaction assays, and lineage-appropriate physiological experiments will be needed to connect the evolutionary patterns reported here to NO signaling functions, including any role in synaptic plasticity. The appropriate interpretation is therefore that this analysis provides a comparative evolutionary framework and a set of testable hypotheses, rather than a demonstration of conserved pathway operation or of causal contributions to nervous-system evolution.

## Limitations

This study draws on currently available proteomes, which continue to expand in both coverage and taxonomic representation. Accordingly, apparent gene absences should be interpreted with appropriate caution. One important limitation remained that phylogenetic reconstruction and reconciliation could only infer evolutionary history; they could not by themselves establish direct functional roles in cognition or plasticity. Therefore, the neurobiological interpretation should be viewed as a testable hypothesis rather than a completed demonstration. Future work combining developmental expression profiling, neuronal localization, biochemical assays, and electrophysiological analyses will be necessary to link these evolutionary patterns to specific roles in LTP, LTD, and higher-order neural function. Even so, the present findings provide an evolutionary framework for investigating whether diversification of NO-related gene families is associated with differences in nervous-system organization and function.

## Conclusion

This comparative phylogenetic analysis shows that putative components of the nitric oxide (NO) signaling system have an ancient and modular evolutionary distribution across the metazoan lineages sampled. NOS homologs were broadly recovered across the examined bilaterian groups and showed a predominantly speciation-associated history, consistent with conservation of an

NO-producing core. In contrast, ASS1, ASL, ARG1, ARG2, PDE5A, GUCY1A1, and GUCY1B1 exhibited more heterogeneous histories, including inferred duplications, divergence, and lineage-specific presence or absence, indicating that substrate-supply, regulatory, and receptor-associated components did not evolve uniformly. Together, these patterns support an evolutionary model in which a comparatively conserved NO-producing component is accompanied by more dynamic diversification of associated pathway gene families.

Because NO–cGMP signaling is implicated in synaptic modulation and plasticity in established experimental systems (Feil and Kleppisch, 2008; Bon and Garthwaite, 2003), the phylogenetic framework developed here provides a basis for examining how these gene families are deployed in metazoan nervous systems. However, comparative proteome and phylogenetic analyses alone cannot establish protein function, pathway activity, neuronal localization, or roles in LTP/LTD. Apparent gene absences should also be interpreted cautiously until supported by improved genome assemblies, annotation, and targeted genomic validation.

### Novelty

The novelty of this work is that it does not examine only one part of the NO pathway. Instead, it connects NO synthesis, arginine recycling, NO regulation, and sGC signaling in the same comparative phylogenetic analysis, and it adds gene-tree/species-tree reconciliation to separate conserved ancestry from lineage-specific change. That makes the study broader than earlier single-gene or single-module studies(Andreakis et al., 2011b; Fitzpatrick et al., 2006; González-Domenech and Muñoz-Chápuli, 2010a; Moroz et al., 2023, 2020b).

## Declarations

### Ethics approval and consent to participate

Not applicable. This study was entirely computational and used only publicly available proteomic and protein sequence data; no human participants, animals, or identifiable private data were involved, so ethics approval and consent to participate were not required.

### Consent for publication

Both authors have read and approved the final manuscript and consent to its submission and publication.

### Availability of data and materials

The supplementary information associated with this manuscript is publicly available. The supplementary materials, including detailed datasets, code, additional information, and analyses, can be accessed via the following figshare doi: https://doi.org/10.6084/m9.figshare.32386266

These materials are provided to ensure transparency, and to facilitate further exploration of the results presented in this study.

## Acknowledgement

We further thank the Central University of Himachal Pradesh (CUHP) for providing computational infrastructure and financial support in the form of a non-NET fellowship.

## Author’s contributions

MK conceptualized the work and AT carried out the research. The results were analyzed and manuscript written by AT and MK.

